# Assignment of the somatic A/B compartments to chromatin domains in giant transcriptionally active lampbrush chromosomes

**DOI:** 10.1101/2023.03.14.532542

**Authors:** Alla Krasikova, Tatiana Kulikova, Juan Sebastian Rodriguez Ramos, Antonina Maslova

**Affiliations:** Saint-Petersburg State University, Saint-Petersburg, Russia

**Keywords:** A/B compartments, chicken genome, chicken karyotype, chromomere, chromomere-loop complex, FISH-mapping, Hi-C, hypertranscription, lampbrush chromosomes, meiotic chromosomes, oogenesis, oocyte nucleus, transcription loops

## Abstract

The three-dimensional configuration of the eukaryotic genome is an emerging area of research. Chromosome conformation capture outlined genome segregation into large scale A and B compartments corresponding mainly to transcriptionally active and repressive chromatin. It remains unknown how the compartmentalization of the genome changes in growing oocytes of animals with hypertranscriptional type of oogenesis. In this type of oogenesis, highly elongated chromosomes, called lampbrush chromosomes, acquire a characteristic chromomere-loop appearance, representing one of the classical model systems for studying the structural and functional organization of chromatin domains. Here, we compared the distribution of A/B compartments in chicken somatic cells with chromatin domains in lampbrush chromosomes. We found that in lampbrush chromosomes, the extended chromatin domains, restricted by compartment boundaries in somatic cells, disintegrate into individual chromomeres. Next, we performed FISH-mapping of the genomic loci, which belong to A or B chromatin compartments as well as to A/B compartment transition regions in embryonic fibroblasts on isolated lampbrush chromosomes. We established, that in chicken lampbrush chromosomes, clusters of dense compact chromomeres bearing short lateral loops and enriched with repressive epigenetic modifications generally correspond to constitutive B compartments in somatic cells. These results suggest that gene-poor regions tend to be packed into chromomeres. Clusters of small loose chromomeres with relatively long lateral loops show no obvious correspondence with either A or B compartment identity. Some genes belonging to facultative B (sub-) compartments can be tissue-specifically transcribed during oogenesis, forming distinct lateral loops.

## Introduction

Nowadays, the three-dimensional chromatin architecture and the mechanisms of its maintenance and reorganization is a rapidly developing area of genome biology [1]. With the advent of chromosome conformation capture approaches, the most widely used of which is the genome-wide Hi-C method, chromatin domains such as A/B compartments, topologically associating domains (TADs), and loop domains have been described [2–5]. A and B compartments are identified from the Hi-C long-range contact frequency matrix by the “checkerboard” pattern and represent large-scale chromatin domains corresponding predominantly to open and closed chromatin [2, 6]. Chromatin visualization confirmed the appearance of active and inactive nuclear compartments, which form two interacting networks in the interphase nucleus and generally match to A and B compartments [7–9]. Moreover, A/B compartment domains may be predicted from the histone modifications profile [10, 11]. A/B compartments have been detected in various organisms, including humans, Drosophila, plants, and recently in the domestic chicken [3, 12–15]. A compartments often correspond to transcriptionally active chromatin and enriched with epigenetic markers of an open chromatin such as H3K36Me3, H3K4me1, H3K27ac, H3K79me2 [3, 16, 17]. B compartments, on the other hand, are usually gene-poor, compact, and contain histone markers of a silent chromatin, such as H3K27me3, H3K9Me3 [18]. In mammals, compartment domains can be subdivided into subcomparments according to epigenetic and transcriptional profile [3, 19]. In contrast to TADs, (sub-) compartment status and boundaries change during cell differentiation that correlates with changes in gene expression profiles [18, 20, 21]. At the same time, there are (sub-) compartments with important housekeeping functions that do not switch their status and therefore can be called constitutive [20].

Several studies indicate that in mammals, conventional compartments are present in early oocytes but become weaker during oocyte maturation and then disappear in maternal zygotic chromatin [22, 23]. Here we aimed to investigate how the compartmentalization of the genome changes in growing oocytes in animals with hypertranscriptional type of oogenesis. In this type of oogenesis, high transcriptional output leads to appearance of extremely elongated lampbrush chromosomes with a typical chromomere-loop structure. Lampbrush chromosomes are found in diplotene stage oocytes of all vertebrates with the exception of mammals [24–28]. At this stage of oogenesis chromosomes nearly lack any *trans*-chromosomal interactions except for the formation of bivalents between homologous chromosomes. In each individual lampbrush chromosome, stable globular chromatin domains, the chromomeres, can be seen along the whole length of the chromosome by conventional light microscopy [29, 30]. Lateral loops emerging from chromomeres are represented by strongly decondensed, actively transcribed chromatin and are visible due to the dense packing of nascent RNA with RNA-binding proteins [31–33]. In birds, lampbrush chromomeres, despite their primarily invariant pattern, are heterogeneous in terms of chromatin compactness, the presence and length of lateral loops, epigenetic modifications, genomic composition, and the abundance of repeats [30, 34–38]. FISH mapping of genomic loci belonging to somatic chromatin domains has not enabled to establish an unambiguous correlation between interphase TADs and lampbrush chromomere-loop complexes [39].

Lampbrush chromosomes are a convenient object for chromatin mapping and have been studied in detail in a number of model organisms [40–42]. However, the correspondence between the chromomere-loop complexes of meiotic lampbrush chromosomes and chromatin compartment identity in the interphase nucleus has not been investigated. We decided to search for regularities in the correspondence between chromatin domains in lampbrush chromosomes of the domestic chicken (*Gallus gallus domesticus*) and A/B compartments in somatic cells using available genome-wide data. To achieve this goal, we set out to compare the pattern of lampbrush chromomeres with A/B compartment distribution along the chromosomes and to map genomic regions belonging to A or B compartments in chicken embryonic fibroblasts on lampbrush chromosomes by FISH. To our knowledge, this is the first study indicating a correlation between the A/B compartments in somatic interphase nucleus and chromatin segments in giant lampbrush chromosomes from diplotene stage oocytes.

## Methods

### A/B compartment distribution analysis

A/B compartments are identified by applying principal component analysis (PCA or eigenvalue decomposition) to the distance-normalized Hi-C interaction frequency matrix for a certain cell type or tissue at a relatively low resolution (usually, bin values are from several tens of thousands kilobases to several megabases). The Pearson correlation matrix is further used to calculate eigenvector for each matrix bin. The type of the compartment (A or B) is generally inferred from the sign of the eigenvector: usually regions with positive values are denoted as “A” compartments, while regions with negative values are denoted as “B” compartments [2]. However, in some cases, additional analysis of GC content or gene density along the particular chromosome being analyzed may be required to correct the eigenvector sign for better prediction of the type of compartment [43]. It has been convincingly shown that GC content and gene density positively correlate with A-compartments and negatively correlate with B compartments [44, 45]. A number of computational program tools have now been developed to generate an A/B compartment profile from the observed/expected Pearson correlation matrix, some of which are embedded in pipelines for processing raw Hi-C datasets, such as *cooltools* (https://open2c.github.io/) and *Juicer tools* ([46]; https://github.com/aidenlab/juicer).

In our study we used publicly available A/B compartment profiles obtained for different chicken cell types: chicken embryonic fibroblasts (CEF) and chicken erythrocytes (RBC) [15], HD3 erythroblasts [47], lymphoblastoid DT40 cells [48], liver cells [49], and chicken small white follicle (SWF) granulosa cells [50]. Since a galGal5 version of the chicken genome assembly was used to generate Hi-C dataset for most cell types except SWF cells, we used this chicken genome assembly to visualize A/B compartment profiles in the Integrative Genomics Viewer (IGV) [51]. Compartment coordinates for SWF cells were remapped to GalGal5 genome assembly using the NCBI remapping service (https://www.ncbi.nlm.nih.gov/genome/tools/remap).

### Labeling the BAC probes by nick-translation

Probes were prepared from the chicken BAC-clone library CHORI-261 (https://bacpacresources.org/chicken261.htm). DNA was extracted from *E*.*coli* night cultures by standard alkaline-lysis protocol according to BAC manufacturer instructions. BAC DNA was labeled by nick-translation [52] at 16°C for 2h, using DNA polymerase I/DNAse I enzyme mix (ThermoFisher Scientific) and either biotin-11-dUTP (Lumiprobe), digoxigenin-11-dUTP (Jena Bioscience) or aminoallyl-dUTP-ATTO-647N (Jena Bioscience). Labeled probes were precipitated and dissolved at 20-30 ng/µl in hybridization mixture, containing 50% formamide, 10% dextran sulfate, 2×SSC and 50× excess of salmon sperm DNA (ThermoFisher Scientific). Since some BAC-clone based probes from the GGA1 gave unspecific hybridization signal on other chromosomes in metaphase plates, in case of these BAC clone combinations, 20× excess of chicken Cot5 DNA prepared by S1-nuclease digestion was added to the hybridization mixture [53]. The probes were preannealed at 37°C for 1h after denaturation before mounting on slides.

### Preparation of lampbrush chromosomes

Lampbrush chromosomes were isolated from growing chicken oocytes according to the earlier described procedure [54, 55] (https://projects.exeter.ac.uk/lampbrush/protocols.htm) under stereomicroscope Leica S9D or M165C (Leica Microsystems). Lampbrush chromosomes were spread by 3000 rpm centrifugation for 30 min at 4°C, fixed in 2% PFA in PBS (1.47 mM KH2PO4, 4.29 mM Na2HPO4, 137 mM NaCl 2.68 mM KCl) for 30 min, dehydrated in ethanol series and air dried. Chickens were handled according to the approval #131-04-6 from 25.03.2019 of the Ethics committee for Animal Research of St. Petersburg State University.

***FISH on lampbrush chromosomes***

DNA/DNA+RNA and DNA/RNA FISH protocols were applied to lampbrush chromosome preparations [56, 57]. Lampbrush chromosomes were denatured in 70% formamide at 70°C for 10-15 min, dehydrated in ice-cold ethanol series and air dried. Hybridization mixtures were denatured at 95°C for 10 min and transferred on ice. 1-5 μl drops of hybridization mixture were applied to lampbrush chromosome preparations covered with coverslip and sealed with rubber cement. Hybridization lasted overnight at 37°C. Post-hybridization washes were in 3 changes of 0.2×SCC at 60°C and 2 changes of 2×2CC at 45°C. Biotin- and digoxigenin-labeled DNA-probes were detected with Alexa488-conjugated streptavidin and Cy3-conjugated anti-digoxigenin antibody correspondingly.

### Microscopy and image analysis

Chromosomes were analyzed with epifluorescence microscope Leica DM4000 (Leica Microsystems), images were acquired with CCD camera (1.3 Mp resolution). DAPI-stained chromosomes were imaged before the FISH procedure, and processed with ImageJ/Fiji multicolor look-up table ‘Fire’ to improve the visibility of brightness heterogeneity along the lampbrush chromosome axis. Schematic drawings of the FISH-mapping of the selected genomic regions on lampbrush chromatin domains represent an approximation of at least 6 images for each chromosomal region.

## Results

### Lampbrush chromosome chromomeric pattern vs. distribution of A/B chromatin compartments

First, we sought to compare the pattern of lampbrush chromomeres with the distribution of A/B compartments along the chromosomes. A/B compartments in interphase nuclei of chicken embryonic fibroblasts (CEF) were recently annotated in Hi-C long-range chromatin interaction maps [15]. Chicken macrochromosomes were characterized by multiple switches between A and B compartments. Here, we used the Integrative Genomics Viewer (IGV) to visualize the distribution of A/B chromatin compartments in CEF and other chicken cell types. Cytological maps of chicken lampbrush chromosomes with a DAPI staining pattern have been developed previously [58–60]. In order to compare the distribution of A/B compartments along the GGA1-GGA7 chromosomes in somatic cells with the distribution of chromomeres on the corresponding lampbrush chromosomes, we used coordinates of previously mapped BAC clones [39, 59–61], genomic positions of centromeres and marker structures such as globular loops [35, 62], as well as the coordinates of individual chromomeres established by microdissection followed by sequencing [35, 37, 38]. The coordinates of the selected markers were refined for the version 5 of the domestic chicken genome (galGal5) (**Additional Table 1**). The results of comparing the distribution of A/B compartments in somatic cells with the pattern of chromomeres along the lampbrush chromosome axes in hen are presented in **Figures 1-5** and **Additional Figure 1**.

**Figure 1.**
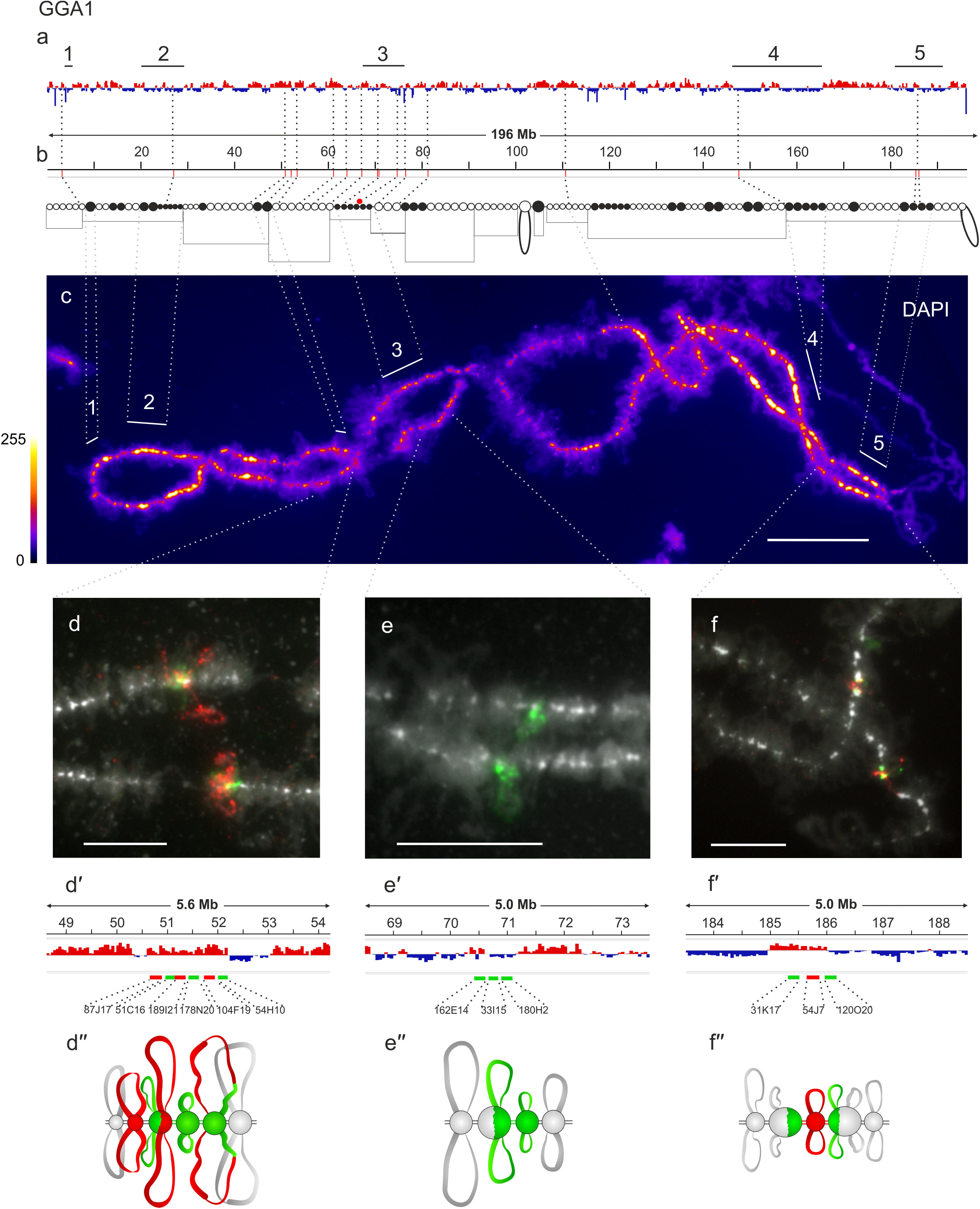
Alignment of interphase genome A/B compartments along chicken chromosome 1 with chromomeric pattern of the corresponding lampbrush chromosome. **a** – distribution of A (red) and B (dark blue) compartments along the chicken chromosome 1 (GGA1) in embryonic fibroblasts viewed by Integrative Genomics Viewer (IGV) (according to [15]); **b** – cytological map of chicken lampbrush chromosome 1 depicting DAPI-staining pattern of chromomeres and relative contour length of lateral loops, black circles – dense chromomeres brightly stained with DAPI (according to [59, 60]). Dotted lines on **a, b** connect the genomic positions of the selected BAC-clones (**Additional Table 1**) with their positions on the cytological map; **c** – lampbrush chromosome 1 stained with DAPI, pixel intensities displayed with multicolored ImageJ look-up table, numbered lines on **a** and **c** indicate positions of chromomere clusters brightly stained with DAPI; **d, e, f** – DNA+RNA-FISH with BAC-clone based DNA-probes (**Additional Table 2**) covering the genomic regions 50-52 Mb (**d**), 70-71 Mb (**e**), and 185-186 Mb (**f**) on chicken lampbrush chromosome 1; dotted lines from **c** to **d**-**f** indicate chromosomal positions of the regions on microphotographs; **d**′-**f**′ – the positions of the mapped BAC-clones relative to the somatic A/B compartments; **d**′′-**f**′′ – schematic drawings of the FISH-mapping of the selected genomic regions on lampbrush chromatin domains; colors correspond to the colors of the labeled DNA-probes on the FISH images. Scale bar: **c** – 20 μm, **d**-**f** – 10 μm.

GGA1 lampbrush chromosome visually differs from the other macrochromosomes by its long length, the absence of any pronounced marker structures, with the exception of the terminal loops (telomere bow-like loops, TBLs) and one marker loop (proximal boundary of axial bar bearing no loops, PBL11) [58, 59, 63]. In lampbrush chromosome 1 several clusters of dense compact chromomeres can be observed (**Figure 1 a-c**). Except for the pericentromeric cluster of chromomeres, all other clusters of prominent DAPI-positive chromomeres (clusters # 1, 2, 4, 5, as well as not numbered smaller regions) correspond to B compartments of different lengths in chicken embryonic fibroblasts.

Lampbrush chromosome GGA2 is characterized by the presence of the so-called “spaghetti’’ marker on the short arm and so-called lumpy loops (LLs) on the long arm [55]. DAPI-positive chromomeres are evenly distributed along the entire length of chromosome 2, with the large clusters in the centromeric region and on the long arm. The locus of “spaghetti marker” formation, the coordinates of which were established earlier [35], corresponds to the A compartment, whereas the cluster of prominent compact chromomeres, in which LLs form [64], corresponds to the B compartment in embryonic fibroblasts (**Figure 2 a-c**). The region of the extended cluster of dense compact chromomeres (cluster #5), the boundaries of which can be tentatively determined using previously mapped BAC clones [59, 60], corresponds to the large B compartment in embryonic fibroblasts (**Figure 2 a-c**).

**Figure 2.**
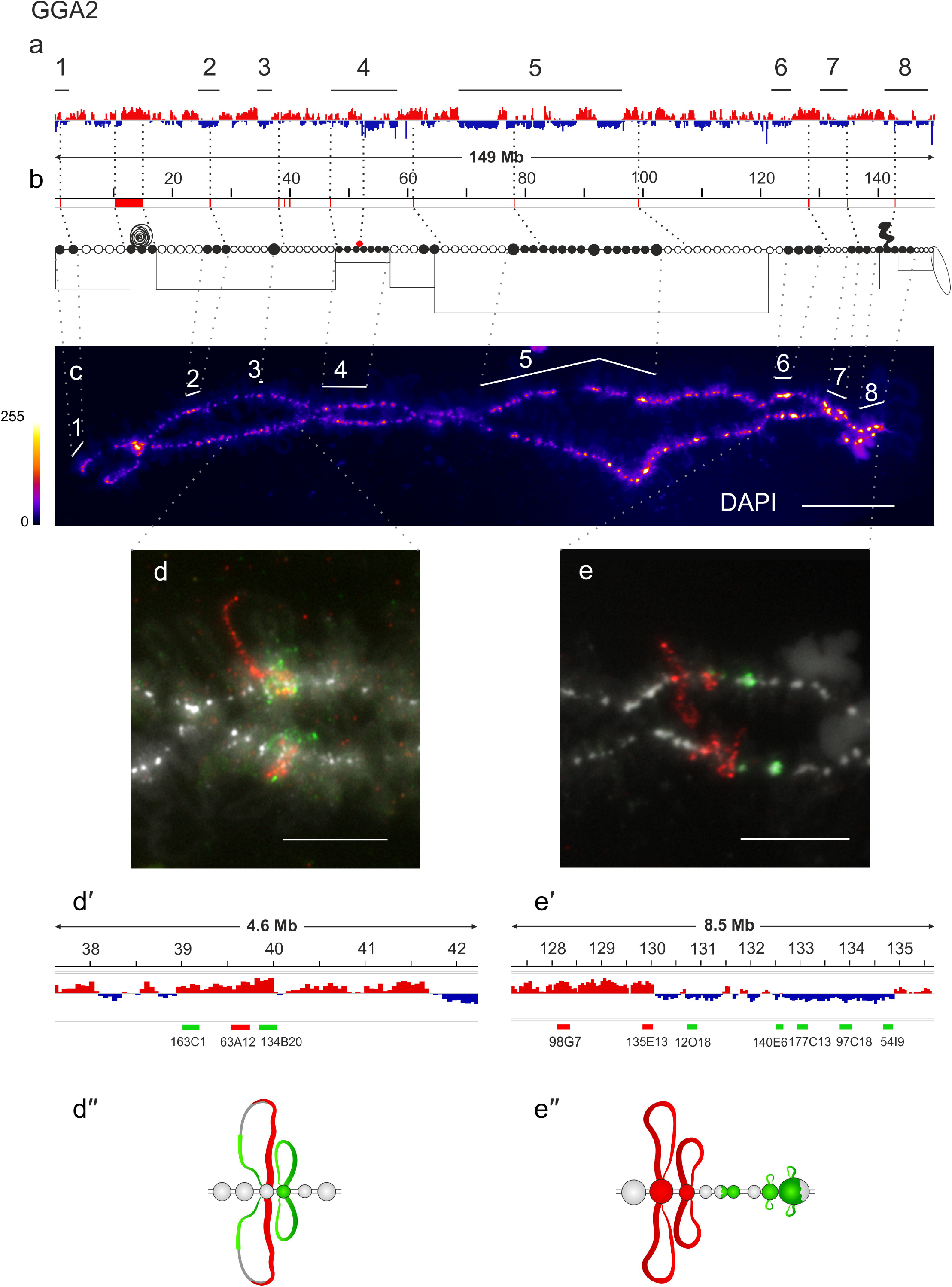
Alignment of interphase genome A/B compartments along chicken chromosome 2 with chromomeric pattern of the corresponding lampbrush chromosome. **a** – distribution of A (red) and B (dark blue) compartments along the chicken chromosome 2 (GGA2) in embryonic fibroblasts viewed by Integrative Genomics Viewer (IGV) (according to [15]); **b** – cytological map of chicken lampbrush chromosome 2 depicting DAPI-staining pattern of chromomeres and relative contour length of lateral loops, black circles – dense chromomeres brightly stained with DAPI (according to [59, 60]). Dotted lines on **a, b** connect the genomic positions of the selected BAC-clones and chromosomal marker structures (**Additional Table 1**) with their positions on the cytological map; **c** – lampbrush chromosome 2 stained with DAPI, pixel intensities displayed with multicolored ImageJ look-up table, numbered lines on **a** and **c** indicate positions of chromomere clusters brightly stained with DAPI; **d, e** – DNA+RNA-FISH with BAC-clone based DNA-probes (**Additional Table 2**) covering the genomic regions 39-40 Mb (**d**), 128-135 Mb (**e**) on chicken lampbrush chromosome 2; dotted lines from **c** to **d** and **e** indicate chromosomal positions of the regions on microphotographs; **d**′, **e**′ – the positions of the mapped BAC-clones relative to the somatic A/B compartments; **d**′′, **e**′′ – schematic drawings of the FISH-mapping of the selected genomic regions on lampbrush chromatin domains; colors correspond to the colors of the labeled DNA-probes on the FISH images. Scale bar: **c** – 20 μm, **d, e** – 10 μm.

The GGA3 lampbrush chromosome is well recognized due to the characteristic asymmetric length of the lateral loops, with a higher average loop length in the region of the long arm closer to the centromere, giving noticeable “fuzziness” in the proximal part of the long arm, whereas the distal part of the long arm is represented mainly by DAPI-positive chromomeres with short lateral loops [58, 59]. These features are well reflected in the distribution of epigenetic modifications: repressive chromatin markers H3K9me3 and 5-methyl-cytosine (5mC) predominate in the clusters of chromomeres in the second half of the long arm [38]. Extended clusters of large chromomeres in this region of lampbrush chromosome 3 generally correspond to the somatic B compartments (**Additional Figure 1 a-a**′′). For instance, the region around the LL marker of chromosome 3, which was mapped before [35], demonstrated the correspondence between 5-6 dense DAPI-positive chromomeres and homogenous B compartment in chicken embryonic fibroblasts.

Lampbrush chromosome GGA4, in turn, has well-defined terminal giant loops (TGLs) recently renamed GITERA (Giant Terminal RNP aggregates) [65], two clusters of DAPI-positive chromomeres, one in the short arm and one in the centromeric region, corresponding to B compartments in embryonic fibroblasts (**Figure 3 a-c**). Chicken lampbrush chromosome 4 is the most completely mapped, both by FISH with BAC clone based probes [39, 59] and by microdissection of individual chromomeres followed by their sequencing [37]. In the proximal part of the long arm of the chromosome 4 is an extended cluster of small DAPI-positive chromomeres (cluster #3) that are depleted in acetylated histone H4 (H4Ac) and H3K9me3 [38]. This region of GGA4 corresponds to intermingled A/B compartments in chicken embryonic fibroblasts with the predominance of B compartment (**Figure 3 a-c**).

**Figure 3.**
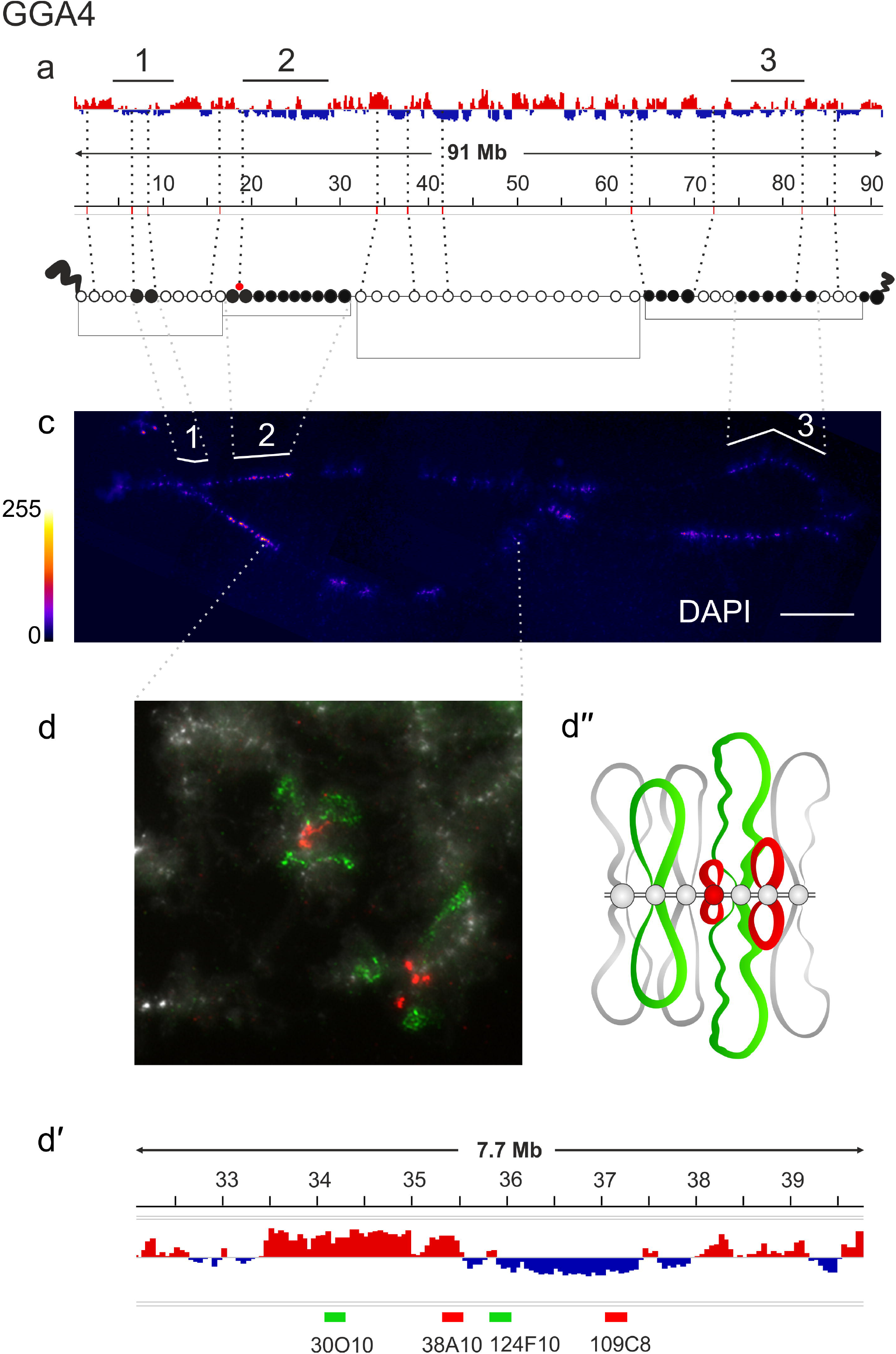
Alignment of interphase genome A/B compartments along chicken chromosome 4 with chromomeric pattern of the corresponding lampbrush chromosome. **a** – distribution of A (red) and B (dark blue) compartments along the chicken chromosome 4 (GGA4) in embryonic fibroblasts viewed by Integrative Genomics Viewer (IGV) (according to [15]); **b** – cytological map of chicken lampbrush chromosome 4 depicting DAPI-staining pattern of chromomeres and relative contour length of lateral loops, black circles – dense chromomeres brightly stained with DAPI (according to [59]). Dotted lines on **a, b** connect the genomic positions of the selected BAC-clones (**Additional Table 1**) with their positions on the cytological map; **c** – lampbrush chromosome 4 stained with DAPI, pixel intensities displayed with multicolored ImageJ look-up table, numbered lines on **a** and **c** indicate positions of chromomere clusters brightly stained with DAPI; **d** – DNA+RNA-FISH with BAC-clone based DNA-probes (**Additional Table 2**) covering the genomic region 34-37 Mb (**d**) on chicken lampbrush chromosome 4; dotted lines from **c** to **d** indicate chromosomal position of the region on microphotographs; **d**′ – the positions of the mapped BAC-clones relative to the somatic A/B compartments; **d**′′ – schematic drawing of the FISH-mapping of the selected genomic region on lampbrush chromatin domains; colors correspond to the colors of the labeled DNA-probes on the FISH image. Scale bar: **c** – 20 μm, **d** – 10 μm.

GGA5 lampbrush chromosome is characterized by several clusters of larger and more globular chromomeres including a cluster without prominent lateral loops in the pericentromeric region [66]. Four of these clusters correlate with the B compartments present at chicken interphase genome (**Additional Figure 1 b-b**′′).

GGA6 in a lampbrush configuration can be divided into two parts – with more compact larger chromomeres and relatively short lateral loops and with tiny chromomeres and longer transcription loops [59]. Prominent B compartment can be assigned to the first part of lampbrush chromosome 6 with higher chromatin compaction (cluster #1) (**Additional Figure 1 c-c**′′).

The GGA7 midichromosome, like the other midichromosomes, is characterized by small DAPI-negative chromomeres from which long lateral loops emerge. Based on the analysis of DAPI staining and counting of the average number of chromomeres, we drew an approximate cytological map of the chromomere pattern for GGA7 (**Figure 4 a-c**). Clusters of DAPI-positive chromomeres are found in the centromeric region, the position of which was previously established [62], and in the distal region of the long arm. B compartments are detected in the genomic region corresponding to these clusters, but further FISH mapping is required to verify their correspondence to the DAPI-positive chromomeres.

**Figure 4.**
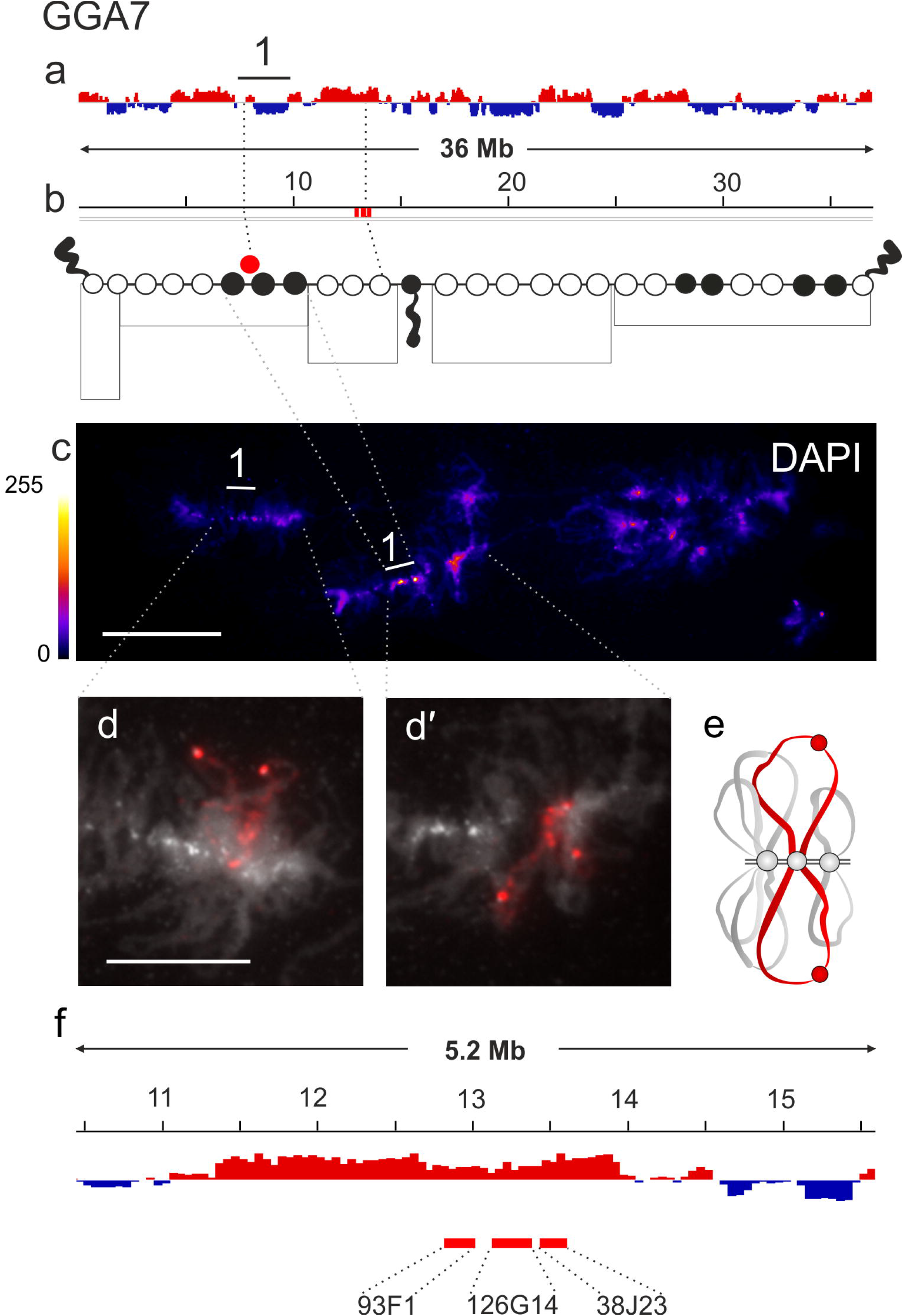
Alignment of interphase genome A/B compartments along chicken chromosome 7 with chromomeric pattern of the corresponding lampbrush chromosome. **a** – distribution of A (red) and B (dark blue) compartments along the chicken chromosome 7 (GGA7) in embryonic fibroblasts viewed by Integrative Genomics Viewer (IGV) (according to [15]); **b** – cytological map of chicken lampbrush chromosome 7 depicting DAPI-staining pattern of chromomeres and relative contour length of lateral loops, black circles – dense chromomeres brightly stained with DAPI. Dotted lines on **a, b** connect the genomic positions of the BAC-clones (**Additional Table 1**) with their positions on the cytological map; **c** – lampbrush chromosome 7 stained with DAPI, pixel intensities displayed with multicolored ImageJ look-up table, numbered line on **a** and **c** indicate position of chromomere cluster brightly stained with DAPI; **d** – DNA+RNA-FISH with BAC-clone based DNA-probes (**Additional Table 2**) covering the genomic region 12-14 Mb (**d, d**′) on chicken lampbrush chromosome 7; dotted lines from **c** to **d, d**′ indicate chromosomal positions of the region on microphotographs; **e** – the positions of the mapped BAC-clones relative to the somatic A/B compartments; **f** – schematic drawing of the FISH-mapping of the selected genomic region (**d, d**′) on lampbrush chromatin domains; colors correspond to the colors of the labeled DNA-probes on the FISH images. Scale bar: **c** – 20 μm, **d, d**′ – 10 μm.

In summary, lampbrush chromosome segments characterized by higher chromatin compaction, more globular chromomeres, and lower transcriptional activity generally correlate with the B compartments present at interphase chromatin.

### FISH-mapping of the genomic regions belonging to A or B compartments on chicken lampbrush chromosomes

Next, we decided to turn our attention to high resolution mapping of the genomic regions belonging to A compartments or regions transitional between A and B compartments on lampbrush chromosome preparations. Based on the available Hi-C data, we selected several regions of interest to obtain probes for FISH mapping of A/B compartments present in chicken interphase genome. For the most part (with a few exceptions), we focused on regions, where the compartment sign was uniform at the 1 to 3 Mb length scales, on the largest macrochromosomes (GGA1-4, GGA7) and one microchromosome (GGA14). Depending on the region, from 2 to 6 BAC clones from the CHORI-261 library were selected to cover the A or B compartment (**Additional Table 2**). The DNA probes derived from the BAC clones to the selected regions of interest were mapped on lampbrush chromosomes by 2D-FISH. FISH was performed according to the DNA/DNA+RNA *in situ* hybridization protocol so that DNA-probes also hybridized with the nascent transcripts on the lateral loops, allowing the hybridization signal to be observed along the entire length of the transcription loop [56].

### Mapping genomic regions belonging to A compartments

To map the genomic regions belonging to A compartments we selected the following segments on four chromosomes: 50-52 Mb in GGA1, 39-40 Mb in GGA2, 12-14 Mb in GGA7, and 1-2 Mb in GGA14 (**Additional Table 2**).

Six BAC clones were selected for the 50-52 Mb region of chromosome 1 (**Figure 1 d**′). FISH mapping of the selected BAC clones on lampbrush chromosome 1 shows the presence of chromomere-loop complexes with long lateral loops in the region corresponding to A compartment (**Figure 1 d-d**′′).

BAC clones 163C1, 63A12, and 134B20 were selected to the 39-40 Mb region of chicken chromosome 2 (GGA2) (**Figure 2 d**′). All three probes hybridized with the RNP-matrix of the lateral loops (**Figure 2 d-d**′′).

Three BAC clones 93F1, 126G14, and 38J23 selected to the 12-14 Mb region of chicken chromosome 7 (GGA7) were mapped into a single pair of long lateral loops with two transcriptional units separated by a chromatin nodule (**Figure 4 d-e**). The presence of a chromatin nodule within the lateral loop can be explained by the fact that in the absence of transcribed genes, chromatin tends to persist in a compact, “closed” state, even without being anchored into a nearby chromomere. Chromatin nodules on the lateral loops of lampbrush chromosomes have been described previously at the morphological level as 5mC enriched transcript-free DNP regions with the characteristic nucleosomal organization [67] and by BAC-clone mapping as untranscribed DNA regions [39].

Two BAC clones, 94D13, 168C19 to a 1-2 Mb region of chicken chromosome 14 (GGA14) also belonging to A compartment hybridized with the RNP matrix of two long lateral loops (**Figure 5 c-c**′′).

**Figure 5.**
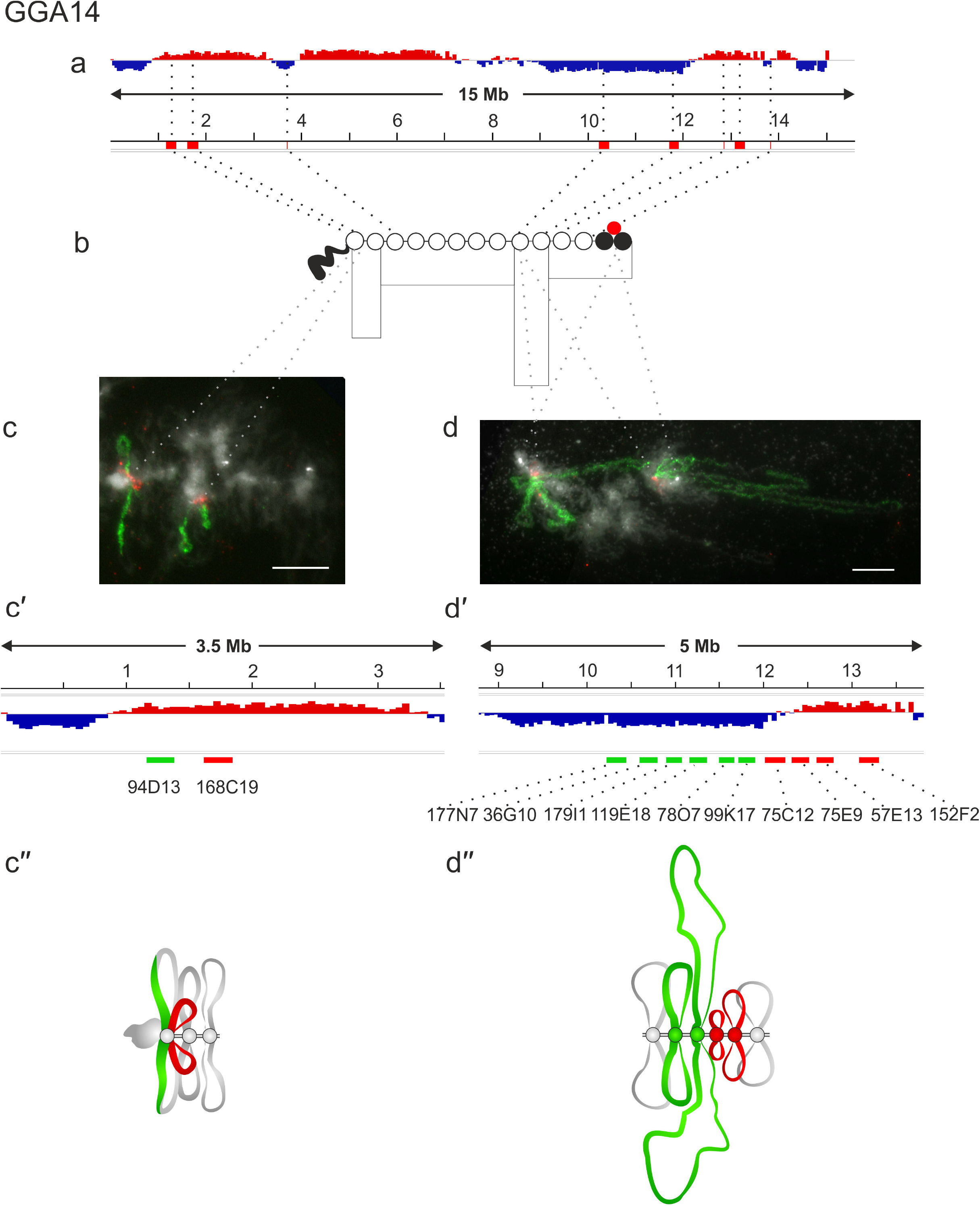
Alignment of interphase genome A/B compartments along chicken chromosome 14 with chromomeric pattern of the corresponding lampbrush chromosome. **a** – distribution of A (red) and B (dark blue) compartments along the chicken chromosome 14 (GGA7) in embryonic fibroblasts viewed by Integrative Genomics Viewer (IGV) (according to [15]); **b** – cytological map of chicken lampbrush chromosome 14 depicting DAPI-staining pattern of chromomeres and relative contour length of lateral loops, black circles – dense chromomeres brightly stained with DAPI (according to [60]). Dotted lines on **a, b** connect the genomic positions of the BAC-clones (**Additional Table 1**) with their positions on the cytological map; **c, d** – DNA+RNA-FISH with BAC-clone based DNA-probes (**Additional Table 2**) covering the genomic regions 1-2 Mb (**c**′) and 10-13 Mb (**d**′) on chicken lampbrush chromosome 14, dotted lines from **b** to **c** and **d** indicate positions of the regions on the cytological map; **c**′, **d**′ – the positions of the mapped BAC-clones relative to the somatic A/B compartments; **c**′′, **d**′′ – schematic drawing of the FISH-mapping of the selected genomic regions (**c, d**) on lampbrush chromatin domains; colors correspond to the colors of the labeled DNA-probes on the FISH images. Scale bar: 10 μm.

### Mapping genomic regions transitional between A and B compartments

We additionally selected BAC clones for A to B compartment transition regions in chicken embryonic fibroblasts for FISH-mapping on lampbrush chromosomes isolated from the oocytes. To this end, 70-71 Mb region in GGA1, 128-135 Mb region in GGA2, 34-37 Mb region in GGA4, and 10-13 Mb region in GGA14 were mapped (**Additional Table 2**).

The 70-71 Mb region of GGA1 is at the border between the chromosomal segment with very long lateral loops and the cluster of centromeric DAPI-positive chromomeres (the so-called ‘centromere bar’). This segment of the lampbrush chromosome 1 corresponds to the interphase chromatin region with multiple switches between A and B compartments (**Figure 1 a-c**). BAC clones 162E14, 33I15, and 180H2 from this region hybridized with two neighboring chromomeres with pairs of short lateral loops (**Figure 1 e-e**′′).

We also selected two differently labeled BAC clone sets to map an extended 128-135 Mb genomic segment of transition from the A to the B compartment on chicken chromosome 2 (GGA2) (**Figure 2 e**′). BAC clones 98G7, 135E13 picked up to the A compartment hybridized into chromomere-loop complexes with long lateral loops, indicating chromatin openness in this region. Next, BAC clones 12O18, 140E6, 177C13, 97C18, and 54I9, matched to the B compartment, hybridized into chromomeres (**Figure 2 e-e**′′). One of these BAC clones also hybridized with a very short lateral loop, which may be related to the transcription of a certain gene. This hybridization pattern fully confirms our observation about the preferential correspondence of A compartments to smaller chromomeres with longer lateral loops and B compartments – to larger chromomeres with shorter lateral loops.

Two other transition genomic regions: the 34-37 Mb segment on chromosome 4 and the 10-13 Mb segment on chromosome 14 demonstrated an exception to the observed tendency. Four BAC clone-based probes, 30O11, 38A10 to the A compartment and 124F10, 109C8 to the B compartment, were selected for the 34-37 Mb region on chromosome 4 (GGA4) (**Figure 3 d**′). All four probes hybridized into the RNP-matrix of four adjacent lateral loops of different sizes regardless of whether they belonged to the A or B compartment (**Figure 3 d-d**′′).

The GGA14 microchromosome contains two clusters of CNM repeats in the centromeric region corresponding to two large DAPI-positive chromomeres at the lampbrush chromosome stage [60] and an additional extended B compartment in the middle of the proximal part of the long arm (**Figure 5 a-b**). At the same time, three BAC clones from this B compartment mapped to three pairs of lateral loops, indicating active transcription of this region at the lampbrush chromosome stage [39]. In particular, the 748 kbp gene *RBFOX1* (RNA Binding Fox-1 Homolog 1) present in this region is transcribed on a very long lateral loop [39] but is silent in chicken embryonic fibroblasts.

Here we selected 9 additional and one previously mapped BAC clones for this region of GGA14 – 6 to the B compartment (177N7, 36G10, 179I1, 119E18, 78O7, 99K17) and 4 to the A compartment (75C12, 75E9, 57E13, 152F2) (10-13 Mb region, **Figures 5 d**′). Probes to the B compartment hybridized with the RNP matrix of two long adjacent lateral loops (**Figure 5 d-d**′′). Probes to the A compartment also hybridized with the lateral loops, but despite the abundance of the genes in this region, the size of these loops was much smaller. This may be related both to the different lengths of the transcribed genes and to the intensity of their transcription at this stage of oogenesis. We assume that in the 10-13 Mb region of GGA14, the longest lateral loops are formed in the genomic regions containing long transcribed genes regardless of the compartment status.

### Mapping genomic regions belonging to B compartments

To map the genomic regions belonging to the somatic B compartments, we chose a 185-186 Mb genomic segment that belongs to GGA1 region, dominated by B compartments interspersed with smaller A compartments (cluster #5) (**Figure 1 a**), and selected three neighboring BAC clones 31K17, 54J7, 120O20 (**Figure 1 f**′). BAC clone-based probes 31K17, 54J7, and 120O20 hybridized to three adjacent chromomeres and probe 54J7 also hybridized with a pair of very short lateral loops, emanating from the central chromomere (**Figure 1 f-f**′′). This pattern of hybridization confirms the general correspondence between genomic regions forming B compartment in chicken embryonic fibroblasts and the large clusters of dense chromomeres with short lateral loops in lampbrush meiotic chromosomes.

In addition, the BAC probes that we earlier mapped on chicken lampbrush chromosome 1 to genomic loci from the two neighboring TADs belonging to the compartment B (146-148 Mb region) hybridize with one prominent chromomere, but not with lateral loops (Figure 2 in [39]). Previously, microdissection and sequencing of a large dense chromomere from this region (149-154 Mb) showed that it is enriched with H3K9me3 and highly methylated gene-poor DNA corresponding to the somatic B compartment [35, 38]. In addition, among three BAC clones to the three subTADs from the compartment B (25-28 Mb region) on lampbrush chromosome 2 (**Figure 2 a**-**c**, region 2), two hybridized with chromomeres and one hybridized with the nascent RNA on a pair of lateral loops corresponding to the *TSHD7A* gene (Figure 3 in [39]).

In conclusion, our data indicate that clusters of dense compact chromomeres carrying short lateral loops and enriched with repressive epigenetic modifications overlap with prominent B compartments in somatic cells, reflecting functional compartmentalization of the chicken genome. Next, we wondered if these compartments are constitutive or cell-type specific. Using the data available up to date, we demonstrate that these B compartments are constitutive between several chicken cell types: embryonic fibroblasts (CEF), mature erythrocytes (RBC), erythroblast cell line (HD3), DT40 cell line, granulosa cells of small white follicle (SWF), and liver cells (**Additional Figure 2**).

## Discussion

The question of whether the structure of lampbrush chromosomes is unique or reflects universal principles of genome packaging remains open. In transcriptionally active lampbrush chromosomes, probably due to the strong repulsion of RNP-matrix of lateral loops, chromatin compartments are apparently absent. Indeed, in somatic cells, genomic loci from one particular B compartment preferentially interact with genomic loci form the other B compartments. However, in giant oocyte nucleus, individual chromosomal segments do not reach each other, both between and within lampbrush chromosomes [68, 69]. Moreover, in growing oocytes of adult birds, lampbrush chromosomes do not interact with nuclear lamina or nucleolus and do not form chromocenters, which reduces the likelihood of interchromosomal interactions. Drosophila polytene chromosomes lack compartments due to similar spatial organization [70].

Previously, based on Hi-C chromatin contact maps, the distribution of TADs and A/B compartments in interphase genome was compared with G-banding (according to Giemsa staining) of human metaphase chromosomes [71]. It was found that TADs with high H1.2/H1X ratio belonging to B compartments strongly correlate with AT-rich Giemsa bands. Our data further indicate that the boundaries between chromatin domains belonging to A/B compartments in somatic cells remain boundaries and in lampbrush chromosomes from diplotene stage oocytes. At the same time, genomic regions corresponding to the extended chromatin domains restricted by compartment boundaries in somatic cells disintegrate into individual chromomeres in diplotene oocytes.

In fact, chicken lampbrush chromosomes are characterized by distinct heterogeneity in chromomere density and contour length of lateral loops. Clusters of dense and relatively large chromomeres with short lateral loops alternate with regions of loose and sometimes indiscernible chromomeres with longer lateral loops [58, 59, 72]. It was found that the repressive chromatin modifications (5mC, H3K9me3, H3K27me3, HP1β) are enriched in the clusters of dense compact chromomeres reflecting their functional state [36, 38]. We found that clusters of more globular chromomeres with relatively short lateral loops and thus higher chromatin compaction generally correspond to the constitutive B compartments present in the interphase nucleus of many cell types. These results suggest that gene-poor regions tend to be packed in chromomeres in lampbrush chromosomes. The boundaries of extended clusters of compact chromomeres corresponding to particular B compartments were established precisely according to the mapped genetic markers. Mapping of the transition regions between A and B compartments further supports the observed tendency. We also assume that in the most genomic regions analyzed, A compartments in chicken embryonic fibroblasts tend to correspond to the lampbrush chromosome segments with smaller and less compact chromomeres and long lateral loops, which correlates with their higher transcriptional status, enrichment with histone H4 acetylation, and elongating form of RNA polymerase II. These chromosomal segments have a greater number of less globular chromomeres compared to the segments overlapping with the somatic B compartments. At the same time, smaller chromomeres with relatively long lateral loops show no obvious correspondence with either A or B compartment identity. A more detailed examination of the distribution of different types of subcompartments and functional classes of chromatin in these genomic regions and their relationship to particular lampbrush chromatin domains is essential.

We also realized that there are a number of exceptions to the observed trend. For example, we noticed that transcription of the lengthy RBFOX1 gene during oogenesis leads to the formation of a long lateral loop in the region corresponding to the B compartment in somatic cells. This is interconnected to oocyte specific expression of many maternal genes as shown by RNA-FISH on lampbrush chromosome preparations [39, 57]. Thus, this B-compartment can be regarded as facultative.

The schematic drawing (**Figure 6**) summarizes the established correspondence between chromatin domains belonging to A or B compartments in chicken interphase genome and lampbrush chromosome segments with different chromomere-loop characteristics. The following issues may be addressed in the future. What is the mechanism of chromatin segregation into individual chromomeres in lampbrush meiotic chromosomes? Whether chromomeres represent chromatin that is passively compacted due to transcriptional inactivity or whether chromatin in chromomeres is protected from transcription initiation due to a more closed chromatin structure? It would be also interesting to compare lampbrush chromomeres with chromatin domains and chromatin domain clusters forming chain-like reticular structure recently described in interphase nucleus by quantitative super-resolution and scanning electron microscopy [7, 73].

**Figure 6.**
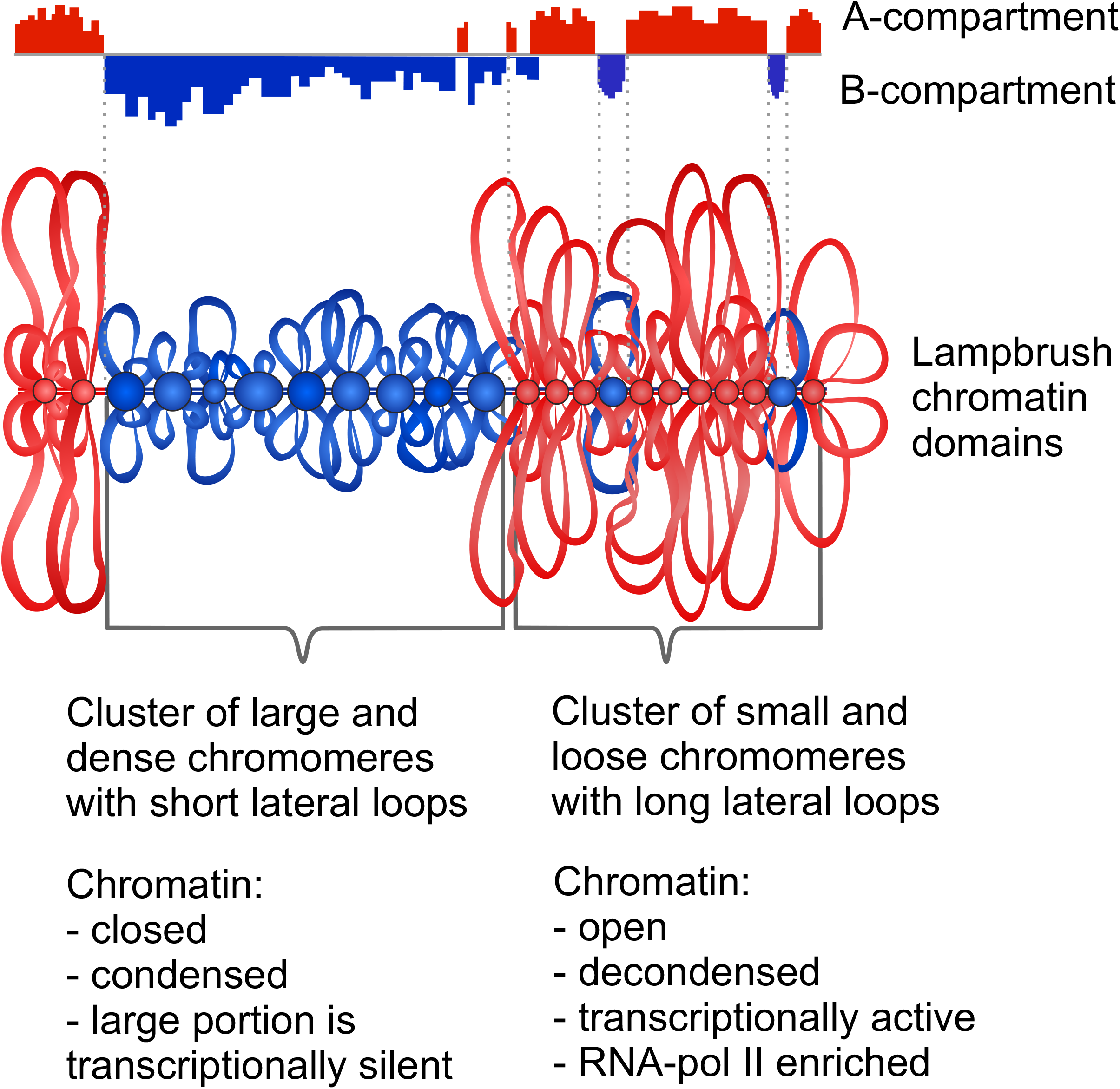
Schematic drawing generalizing the correspondence between A or B compartments present in interphase genome and lampbrush chromosome segments. Chromosomal segments with more globular compact chromomeres and short lateral loops correspond to B compartment (blue), whereas chromosomal segments composed by small loose chromomeres with long transcription loops correspond to A compartment (red).

## Acknowledgements

The study was supported by the RSF grant #19-74-20075. Microscopy was performed using the equipment of the Resource Center “Molecular and Cell Technologies” (Research Park of Saint-Petersburg State University).

## Authors’ contributions

AK conceptualized the study and supervised the project. JS RR, TK and AK compared the distribution of compartments with lampbrush chromomere pattern. AM and AK selected regions of interest and BAC clone-based probes, AM labeled probes by nick-translation. TK and JS RR performed FISH on isolated lampbrush chromosomes and analyzed the micrographs. AK suggested the final scheme. AK and TK wrote the manuscript with a contribution from co-authors. All authors read and approved the final manuscript.

## Additional Material

**Additional Figure 1**. Distribution of A/B compartments along the chicken chromosome 3, 5 and 6 in embryonic fibroblasts compared with the chromomeric pattern in the corresponding lampbrush chromosomes.

**Additional Figure 2**. Examples of constitutive B compartment regions in previously studied chicken cell types.

**Additional Table 1**. Genomic regions mapped on chicken lampbrush chromosomes and marked on the coordinate line on Figures 1-5, Additional Figure 1, according to the chicken genome version 5 (galGal5).

**Additional Table 2**. The list of BAC clones containing fragments of chicken genomic DNA from the CHORI-261 library that were used as DNA-probes for FISH; coordinates are indicated according to the chicken genome version 5 (galGal5).

## Notes

### Competing Interest Statement

The authors have declared no competing interest.

