## Additional Figure for "Assignment of the somatic A/B compartments to chromatin domains in giant transcriptionally active lampbrush chromosomes"

**Additional Material**

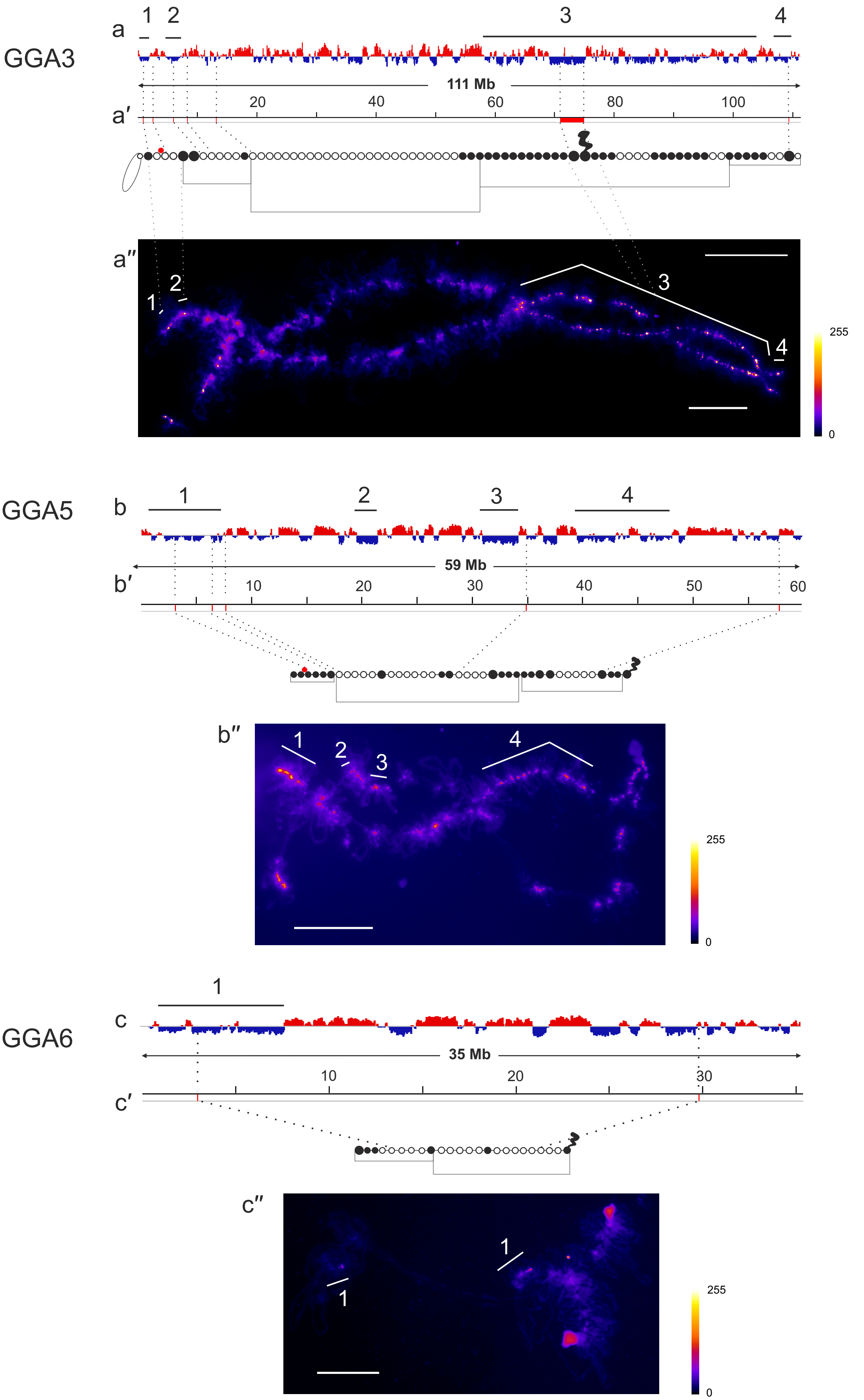

GGA3
a
1
2
3
4
111 Mb
a′
20
40
60
80
100
a″
255
0
GGA5
b
59 Mb
b′
10
20
30
40
50
60
b″
GGA6
c
35 Mb
c′
c″

**Additional Figure 1. Distribution of A/B compartments along the chicken chromosome 3, 5 and 6 in embryonic fibroblasts compared with the chromomeric pattern in the corresponding lampbrush chromosomes.**

Distribution of A (red) and B (dark blue) compartments along the chicken chromosomes: 3 (GGA3) (**a**), 5 (GGA5) (**b**) and 6 (GGA6) (**c**), in embryonic fibroblasts viewed by Integrative Genomics Viewer (IGV) (according to [15]); cytological maps of chicken lampbrush chromosomes 3 (**a′**), 5 (**b′**), 6 (**c′**) depicting DAPI-staining pattern of chromomeres and relative contour length of lateral loops, black circles – dense chromomeres brightly stained with DAPI (according to [59, 60]). Dotted lines connect the genomic positions of the BAC-clones (Additional Table 1) with their positions on the cytological maps. **a″**-**c″** – lampbrush chromosomes 3 (**a″**), 5 (**b″**), 6 (**c″**) stained with DAPI, pixel intensities displayed with multicolored ImageJ look-up table, numbered lines on **a**-**c** and **a″**-**c″** indicate positions of chromomere clusters brightly stained with DAPI. Scale bars: 20 μm.

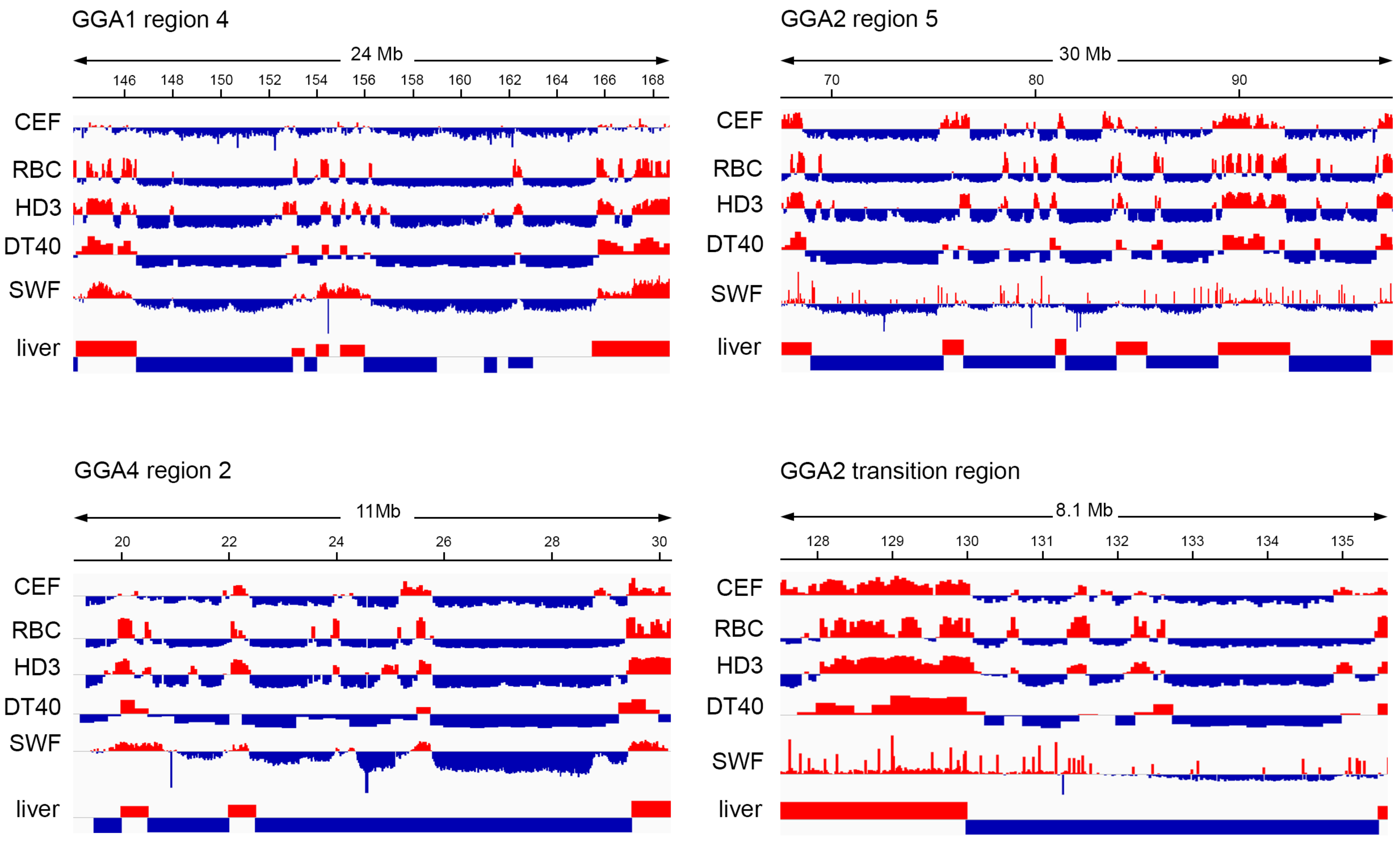

GGA1 region 4
24 Mb
146 148 150 152 154 156 158 160 162 164 166 168
CEF
RBC
HD3
DT40
SWF
liver
GGA2 region 5
30 Mb
70 80 90
CEF
RBC
HD3
DT40
SWF
liver
GGA4 region 2
11Mb
20 22 24 26 28 30
CEF
RBC
HD3
DT40
SWF
liver
GGA2 transition region
8.1 Mb
128 129 130 131 132 133 134 135
CEF
RBC
HD3
DT40
SWF
liver

**Additional Figure 2. Examples of constitutive B compartment regions in previously studied chicken cell types.**

The regions of large constitutive B compartment domains on chromosomes GGA1, GGA2 and GGA4 and the conservative A-to-B compartment transition region on GGA2. The profiles of A (red) and B (blue) compartment for chicken embryonic fibroblasts (CEF, resolution – 50 Kb), chicken erythrocytes (RBC, resolution – 50 Kb), HD3 erythroblasts (resolution – 50 Kb), DT40 cells (resolution – 250 Kb), small white follicle granulosa cells (SWF, resolution – 20 Kb), and liver cells (resolution – 500 Kb) are plotted in the Integrative Genomics Viewer (IGV).

**Additional Table 1**. Genomic regions mapped on chicken lampbrush chromosomes and marked on the coordinate line on **Figures 1-5**, **Additional Figure 1**, according to the chicken genome version 5 (galGal5).

| Chromosome | DNA-probe | Genetic marker | Genomic coordinates (bp) | | Reference |
|---|---|---|---|---|---|
| GGA1 | W030P07 | MCW248 | 3042685 | 3042903 | [59] |
| GGA1 | W030B21 | LEI194 | 26981390 | 26981528 | [59] |
| GGA1 | CH261-87J17 | | 50659355 | 50900102 | *current study* |
| GGA1 | CH261-54H10 | | 52008845 | 52206496 | *current study* |
| GGA1 | WAG31B10 | LEI146 | 53274224 | 53274474 | [59, 60] |
| GGA1 | WAG13E20 | GCT0049 | 60975037 | 60975254 | [60] |
| GGA1 | WAG69C11 | MCW007 | 63858174 | 63858471 | [60] |
| GGA1 | WAG67J15 | LDHB | 67084357 | 67094049 | [60] |
| GGA1 | CH261-162E14 | | 70424086 | 70619427 | *current study* |
| GGA1 | CH261-33I15 | | 70675137 | 70847856 | *current study* |
| GGA1 | WAG53E23 | LEI0071 | 76397821 | 76397841 | [60] |
| GGA1 | WAG25G16 | LEI101 | 81210437 | 81210594 | [60] |
| GGA1 | P1H9 | GCT13 | 110607927 | 110608236 | [59] |
| GGA1 | CH261-191J12 | | 147563186 | 147752805 | [39] |
| GGA1 | CH261-54J7 | | 185317106 | 185519653 | *current study* |
| GGA1 | CH261-120O20 | | 185970069 | 186181883 | *current study* |
| GGA2 | W055L19 | ADL228 | 830924 | 830943 | [59] |
| GGA2 | Spagetti marker | | 10300000 | 15000000 | [35] |
| GGA2 | CH261-96F4 | | 26339255 | 26564682 | [39] |
| GGA2 | W026B13 | MCW63 | 38085213 | 38085346 | [59] |
| GGA2 | CH261-163C1 | | 39010879 | 39197698 | *current study* |
| GGA2 | CH261-134B20 | | 39844735 | 40041305 | *current study* |
| GGA2 | CH261-182I12 | | 46712094 | 46921822 | [39] |
| GGA2 | W014J06 | MCW358 | 60810746 | 60810764 | [60] |
| GGA2 | P2E4 | GCT23 | 78032760 | 78032777 | [59] |
| GGA2 | W041C02 | LEI147 | 99097193 | 99096954 | [59, 60] |
| GGA2 | CH261-98G7 | | 128115147 | 128367991 | *current study* |
| GGA2 | CH261-54I9 | | 134678478 | 134876251 | *current study* |
| GGA2 | LL2R | | 142822175 | 142893497 | [64] |
| GGA3 | WAG29L12 | MCW0261 | 835733 | 835983 | [60] |
| GGA3 | WAG54M22 | MCW0141 | 2558467 | 2558696 | [60] |
| GGA3 | WAG13D11 | CZ566991 | 5944007 | 5944620 | [60] |
| GGA3 | WAG40J15 | CZ564186 | 8278622 | 8279152 | [60] |
| GGA3 | WAG32A13 | CZ562529 | 13128312 | 13128887 | [60] |
| GGA3 | chromomere #16-7 | | 71000000 | 750000000 | [35] |
| GGA3 | W013L14 | MCW37 | 109303638 | 109303891 | [59] |
| GGA4 | W008H20 | ADL143 | 1431102 | 1431253 | [59] |
| GGA4 | CH261-33C6 | | 6419141 | 6610711 | [39] |
| GGA4 | W023I16 | ADL203 | 8314125 | 8314310 | [59] |
| GGA4 | W125P16 | MCW295 | 16406325 | 16406429 | [59] |
| GGA4 | CH261-30O11 | | 34089061 | 34311128 | *current study* |
| GGA4 | CH261-49F17 | | 37602285 | 37797874 | [39] |
| GGA4 | CH261-47F11 | | 41547668 | 41747405 | [39] |
| GGA4 | CH261-139O9 | | 62817075 | 62995554 | [39] |
| GGA4 | W012C06 | MCW180 | 72199515 | 72199534 | [59] |
| GGA4 | W013E02 | LEI63 | 82180055 | 82179854 | [59] |
| GGA4 | W037E19 | LEI73 | 85792844 | 85793001 | [59] |
| GGA5 | 231H07 | | 3057165 | 3057166 | [66] |
| GGA5 | 429G11 | | 6454339 | 6454340 | [66] |
| GGA5 | W037H20 | MCW263 | 7666396 | 7666785 | [59] |
| GGA5 | W003K18 | MCW210 | 34881129 | 34881612 | [59] |
| GGA5 | W009B13 | ADL298 | 57799040 | 57799394 | [59] |
| GGA6 | W027G19 | LEI192 | 2943525 | 2943846 | [59] |
| GGA6 | W010H24 | ADL142 | 29784025 | 29784326 | [59] |
| GGA7 | CH261-93F1 | | 12820752 | 13020947 | *current study* |
| GGA7 | CH261-126G14 | | 13125892 | 13385143 | *current study* |
| GGA7 | CH261-38J23 | | 13436411 | 13615182 | *current study* |
| GGA14 | CH261-94D13 | | 1164728 | 1385107 | *current study* |
| GGA14 | CH261-168C19 | | 1620727 | 1848725 | *current study* |
| GGA14 | WAG32F10 | | 3695102 | 3695437 | [60] |
| GGA14 | CH261-177N7 | | 10232089 | 10454271 | *current study* |
| GGA14 | CH261-99K17 | | 11716239 | 11911222 | *current study* |
| GGA14 | WAG19G22 | | 12851870 | 12852527 | [60] |
| GGA14 | CH261-152F2 | | 13090203 | 13310067 | *current study* |
| GGA14 | WAG42M3 | | 13831988 | 13832434 | [60] |

**Additional Table 2**. The list of BAC clones containing fragments of chicken genomic DNA from the CHORI-261 library that were used as DNA-probes for FISH, coordinates are indicated according to the chicken genome version 5 (galGal5).

| Chromosome region (Mb) | BAC clone name | Start coordinate (bp) | End coordinate (bp) | Insert lenght (bp) |
|---|---|---|---|---|
| GGA1_50-52 | 87J17 | 50659355 | 50900102 | 240748 |
| GGA1_50-52 | 51C16 | 50965726 | 51148987 | 183262 |
| GGA1_50-52 | 189I21 | 51149051 | 51371768 | 222718 |
| GGA1_50-52 | 178N20 | 51428246 | 51636410 | 208165 |
| GGA1_50-52 | 104F19 | 51731158 | 51950181 | 219024 |
| GGA1_50-52 | 54H10 | 52008845 | 52206496 | 197652 |
| GGA1_70-71 | 162E14 | 70424086 | 70619427 | 195342 |
| GGA1_70-71 | 33I15 | 70675137 | 70847856 | 172720 |
| GGA1_70-71 | 180H2 | 70897644 | 71097854 | 200211 |
| GGA1_185-186 | 31K17 | 185317106 | 185519653 | 202548 |
| GGA1_185-186 | 54J7 | 185647688 | 185877592 | 229905 |
| GGA1_185-186 | 120O20 | 185970069 | 186181883 | 211815 |
| GGA2_39-40 | 163C1 | 39010879 | 39197698 | 186820 |
| GGA2_39-40 | 63A12 | 39544475 | 39746185 | 201711 |
| GGA2_39-40 | 134B20 | 39844735 | 40041305 | 196571 |
| GGA2_128-135 | 98G7 | 128115147 | 128367991 | 252845 |
| GGA2_128-135 | 135E13 | 129835347 | 130046990 | 211644 |
| GGA2_128-135 | 12O18 | 130737070 | 130936225 | 199156 |
| GGA2_128-135 | 140E6 | 132515988 | 132668627 | 152640 |
| GGA2_128-135 | 177C13 | 132943066 | 133156205 | 213140 |
| GGA2_128-135 | 97C18 | 133802274 | 134043084 | 240811 |
| GGA2_128-135 | 54I9 | 134678478 | 134876251 | 197774 |
| GGA4_34-37 | 30O11 | 34089061 | 34311128 | 222068 |
| GGA4_34-37 | 38A10 | 35326117 | 35549083 | 222967 |
| GGA4_34-37 | 124F10 | 35824748 | 36061806 | 237059 |
| GGA4_34-37 | 109C8 | 37052261 | 37282110 | 229850 |
| GGA7_12-14 | 93F1 | 12820752 | 13020947 | 200196 |
| GGA7_12-14 | 126G14 | 13125892 | 13385143 | 259252 |
| GGA7_12-14 | 38J23 | 13436411 | 13615182 | 178772 |
| GGA14_1-2 | 94D13 | 1164728 | 1385107 | 220380 |
| GGA14_1-2 | 168C19 | 1620727 | 1848725 | 227999 |
| GGA14_10-13 | 177N7 | 10232089 | 10454271 | 222183 |
| GGA14_10-13 | 36G10 | 10599836 | 10804688 | 204853 |
| GGA14_10-13 | 179I1 | 10900679 | 11080896 | 180218 |
| GGA14_10-13 | 119E18 | 11168171 | 11366145 | 197975 |
| GGA14_10-13 | 78O7 | 11501758 | 11675622 | 173865 |
| GGA14_10-13 | 99K17 | 11716239 | 11911222 | 194984 |
| GGA14_10-13 | 75C12 | 12023591 | 12250932 | 227342 |
| GGA14_10-13 | 75E9 | 12322420 | 12523919 | 201500 |
| GGA14_10-13 | 57E13 | 12604054 | 12796309 | 192256 |
| GGA14_10-13 | 152F2 | 13090203 | 13310067 | 219865 |
